# High-throughput algorithm predicts F-Type ATP synthase rotor ring stoichiometries of 8 to 27 protomers

**DOI:** 10.1101/2024.02.27.582367

**Authors:** Stepan D. Osipov, Egor V. Zinovev, Arina A. Anuchina, Alexander S. Kuzmin, Andronika V. Minaeva, Yury L. Ryzhykau, Alexey V. Vlasov, Ivan Yu. Gushchin

## Abstract

ATP synthases are large enzymes present in every living cell. They consist of a transmembrane and a soluble domain, each comprising multiple subunits. The transmembrane part contains an oligomeric rotor ring (c-ring), whose stoichiometry defines the ratio between the number of synthesized ATP molecules and the number of ions transported through the membrane. Currently, c-rings of F-Type ATP synthases consisting of 8 to 17 (except 16) subunits have been experimentally demonstrated. Here, we present an easy-to-use high-throughput computational approach based on AlphaFold that allows us to estimate the stoichiometry of all homooligomeric c-rings, whose sequences are present in genomic databases. We validate the approach on the available experimental data, obtaining the correlation as high as 0.94 for the reference data set, and use it to predict the existence of c-rings with stoichiometry varying from 8 to 27. We then conduct molecular dynamics simulations of two c-rings with stoichiometry above 17 to corroborate the machine learning-based predictions. Our work strongly suggests existence of rotor rings with previously undescribed high stoichiometry in natural organisms and highlights the utility of AlphaFold-based approaches for studying homooligomeric proteins.

## Introduction

Adenosine triphosphate (ATP) synthases are crucial enzymes present in every living cell as they convert energy of the transmembrane potential into that of ATP’s chemical bonds (Junge and Nelson 2015; Kühlbrandt 2019; Nirody, Budin and Rangamani 2020; Zubareva *et al*. 2020). According to modern classification, ATP synthases are divided into F-type, found in bacteria, and chloroplasts and mitochondria of eukaryotes, and A-type, found in archaea (Niu *et al*. 2017). ATP synthases are also related to vacuolar-type ATPases (Wang and Rubinstein 2023). Here, we focus on F-type ATP synthases (F-ATP synthases).

F-ATP synthases consist of a transmembrane domain F_O_ and a soluble domain F_1_, each comprising multiple subunits. While one part of ATP synthase remains relatively static during the enzyme operation, another one rotates, being driven by ion flow (H^+^ or Na^+^) that follows the gradient of electrochemical potential across the membrane. The transmembrane part of ATP synthases contains a rotor ring, called c_N_-ring, an oligomer of N subunits of type c. Stoichiometry of the rotor ring N defines the ratio between the number of synthesized ATP molecules and the number of transported ions (3/N), since every complete turn of the c_N_-ring results in synthesis of 3 ATPs from 3 adenosine diphosphates (ADPs). Thus, the ratio 3/N defines one of the main bioenergetic parameters for each living organism.

Currently, a plethora of experimentally obtained high-resolution structures of ATP synthases from different species are available in PDB (Kühlbrandt 2019; Vlasov et al. 2022) as well as of structures of separate transmembrane rotor rings (Meier *et al*. 2009; Pogoryelov *et al*. 2010; Symersky *et al*. 2012; Schulz *et al*. 2013; Matthies *et al*. 2014; Preiss *et al*. 2014, 2015). Each subunit c, a protomer of the rotor ring, consists of two transmembrane α-helices connected by a loop, forming a hairpin-like structure, with the N-terminal helix being on the inner side of the c-ring, and the C-terminal helix on the outer side. The N- and C-termini of subunit c are positioned on one side of the membrane, and the conservative loop contacts the γ-subunit on the other side of the membrane.

c-rings from different species have different stoichiometries (Vlasov et al., 2022), varying from 8 protomers in rotor rings of mammalian mitochondrial ATP synthases (Gu *et al*. 2019; Spikes, Montgomery and Walker 2020) to 15 or 17 subunits in some bacterial species (*Arthrospira platensis* (Pogoryelov *et al*. 2009) and *Burkholderia pseudomallei* (Schulz *et al*. 2017), respectively). In addition, V-type ATPase rings consisting of 10 subunits with 4 α-helices (thus corresponding to c_20_ ring of F-Type ATPases) were reported (Murata *et al*. 2005). Given that c-ring stoichiometry defines the efficiency of ATP synthase, studying its key determinants is of utmost importance. Apparently, c-ring stoichiometry is defined by the amino acids placed at the interfaces between adjacent c subunits, and usage of mutations to alter stoichiometry was previously demonstrated: natural c_11_-ring from *Ilyobacter tartaricus* was transformed into c_<11_, c_12_, c_13_, c_14_, and c_>14_ rings (Pogoryelov et al. 2012); c_13_-ring from *Bacillus pseudofirmus OF4* was transformed into c_12_ rings (Preiss *et al*. 2013); and c_14_ ring from chloroplasts of a plant *Nicotiana tabacum* was transformed into c_15_ ring (Yamamoto *et al*. 2023). Engineered c-rings could be used for modulating organism properties, as demonstrated for example in Yamamoto *et al*. 2023. However, c-ring stoichiometry engineering is still time- and effort-consuming, and lacks a straightforward computational approach that would accurately predict c-ring stoichiometry.

Recently, development of AlphaFold2 provided a highly efficient and reliable machine learning-based approach for protein structure prediction, which revolutionized structural biology (Jumper et al. 2021; Lupas et al. 2021; Akdel et al. 2022). Several other approaches with similar performance have been released soon thereafter (Baek et al. 2021; Chowdhury et al. 2022; Lin et al. 2023). Community quickly realized that the abilities of these approaches go beyond the original objectives (Akdel et al. 2022). In particular, efficient prediction of structures of homo- and hetero-oligomeric assemblies was demonstrated (Akdel et al. 2022; Bryant et al. 2022a; Schweke et al. 2023; Shor and Schneidman-Duhovny 2024). While the available approaches perform well on low to moderate stoichiometry targets, their performance on high stoichiometry homooligomeric proteins, particularly ATP synthase c-rings, hasn’t been tested thoroughly.

In this work, we present an easy-to-use approach, which allows prediction of the stoichiometry of c-rings. The method demonstrates high accuracy of predictions and does not require large computational resources. Consequently, we estimated the stoichiometry of more than one thousand representative c-rings from different species. We predict the existence of previously undescribed high-stoichiometry native rotor rings (up to 27) in a number of species and corroborate the findings by conducting molecular dynamics simulations of exemplary c-rings. We conclude that the existence of ATP synthases with high-stoichiometry c-rings, which might operate at lower-than-expected transmembrane potentials or higher adverse ion concentration gradients, is highly likely and that AlphaFold-based approaches hold a great potential for evaluation of natural diversity of homooligomeric protein complexes.

## Results

Previously, several approaches to AlphaFold2-based stoichiometry prediction were reported that relied on modeling of homooligomers with all possible stoichiometries (Akdel et al. 2022; Schweke et al. 2024; Shor and Schneidman-Duhovny 2024). There, a certain metric was evaluated for each oligomer to determine the most probably stoichiometry: pTM score by Akdel et al. 2022, PAE-based confidence score by Shor and Schneidman-Duhovny 2024, and number of steric clashes by Schweke et al. 2024. Also, reported approaches were tested mostly for low-stoichiometry relatively large rigid proteins. Provided that c-ring stoichiometry could reach at least 20, using such approaches would require obtaining a large number of possible models, making the process computationally expensive and sometimes beyond reach of AlphaFold and ColabFold implementations run on standard workstations or cloud servers. When used for modeling the spinach chloroplast ATP synthase c-ring, AlphaFold2 predicted complete rings for assemblies including 10 to 21 subunits (Supporting Figure 1), whereas correct stoichiometry is 14.

**Figure 1.**
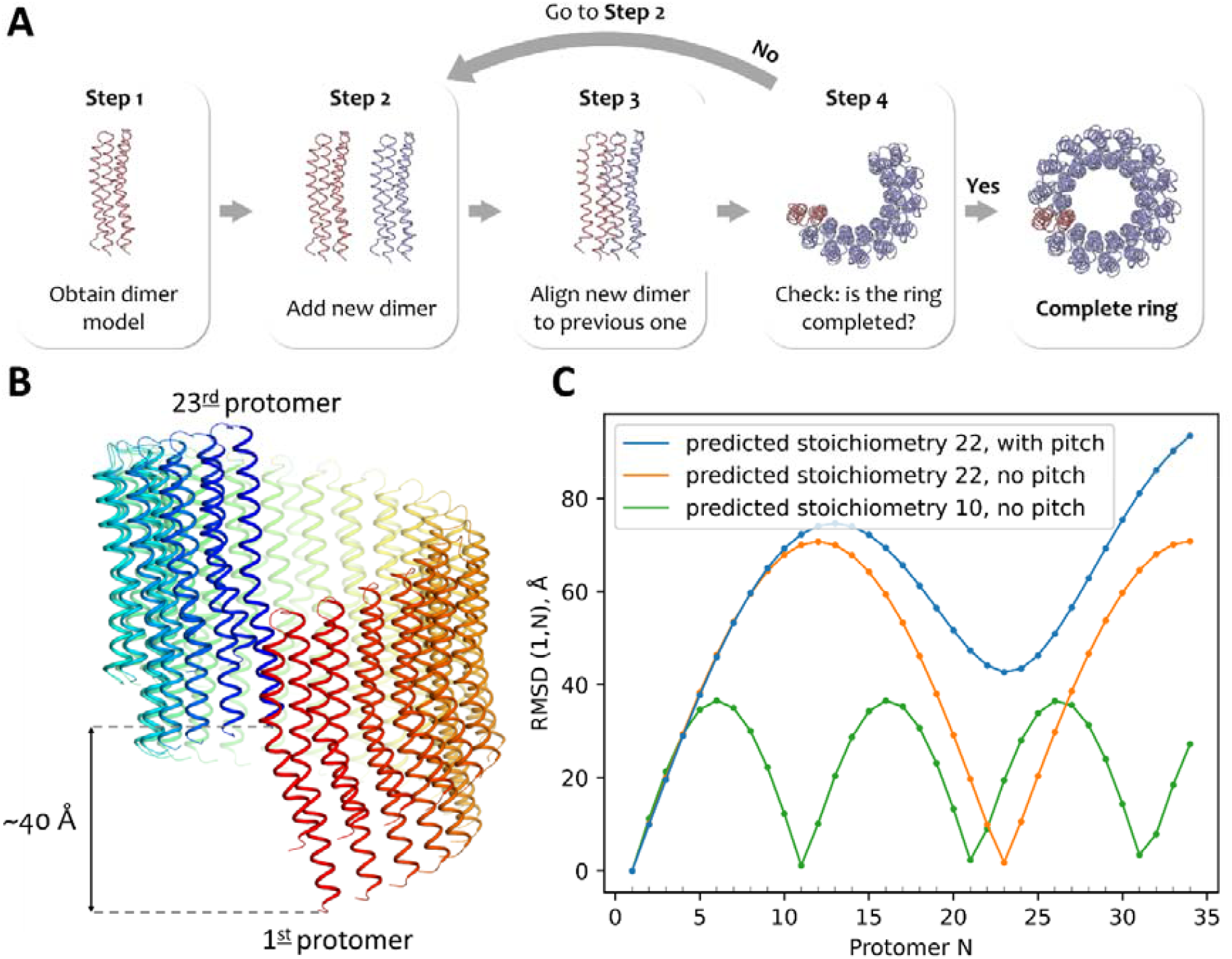
Stoichiometry prediction algorithm. **(A)** Block scheme of the proposed algorithm. Step 1: AlphaFold2-based ColabFold is used to predict the relative position of two protomers, #1 and #2, within a model of a dimer. Step 2: new dimer, which will correspond to protomers #N+1 and #N, is generated by duplication of the original model. Step 3: protomer #N of the new dimer is aligned to the last protomer of the previously generated dimer. Step 4: a check is performed whether the ring is completed; if not, steps 2-4 are repeated. To check whether the ring is completed, root-mean-square deviation (RMSD) between the atom positions of protomer #N+1 and #1, RMSD(1,N+1), is calculated. If RMSD(1,N+1) becomes higher than RMSD(1,N), the stoichiometry is assumed to be N. **(B)** Example of a modeled oligomeric assembly displaying pronounced helical pitch (Uniprot ID A0A972IW59); most of the c subunit oligomer models do not display any pitch. **(C)** Plots showing RMSD(1,N) for three exemplary cases (Uniprot IDs A0A972IW59, A0A1L3GRB0, A0A074LU04). Pronounced troughs correspond to complete circular or helical repeats.

Consequently, we decided to test an alternative approach, where a small oligomer is modeled first to determine relative position of protomers, and then the obtained mutual arrangement is used to extend the assembly aiming for closure of the oligomeric ring (Figure 1). The approach is partially inspired by the DockTrina method developed for docking of trimeric proteins (Popov, Ritchie and Grudinin 2014) and AnAnaS method for analysis of cyclic symmetries (Pagès, Kinzina and Grudinin 2018).

### Development and validation of stoichiometry prediction algorithm

Initially, it was not clear to us, which oligomer should be used preferably as the template for extracting relative positions of two protomers in the complete ring. On one hand, structure prediction is faster for smaller models, so modeling a dimer would be the most time-efficient approach. On the other hand, c subunits of F-ATPases consist of only two α-helices, contacting one neighboring protomer on one side and another protomer on the other side. Modeling a dimer would leave each protomer with an exposed interface, and absence of neighbors could alter the structure of each protomer and thus their mutual positions and the interface between them.

Consequently, we tested the stoichiometry predictions for the reference data set of c-rings with experimentally determined structures (Supporting Table 1) for different interfaces obtained from dimer, trimer and tetramer models. Somewhat surprisingly, we observed very good correlation (0.94) between the experimental and predicted values for interfaces extracted from dimer models (Supporting Table 2). Overall, the algorithm tended to slightly overestimate the stoichiometry (Figure 2), but since it is not clear whether, and how, this trend should be extrapolated for stoichiometries above 17, we decided to analyze the predictions as is, without correcting for this overestimation (but keeping it in mind for prediction-based conclusions).

**Table 1.** ColabFold parameters used to generate dimer models.

|  |  |
| --- | --- |
| **msa_mode** | MMseqs2 (UniRef+Environmental) |
| **num_models** | 5 |
| **num_recycles** | 3 |
| **stop_at_score** | 100 |
| **num_relax** | 0 |
| **relax_max_iterations** | 200 |
| **use_templates** | No |

**Figure 2.**
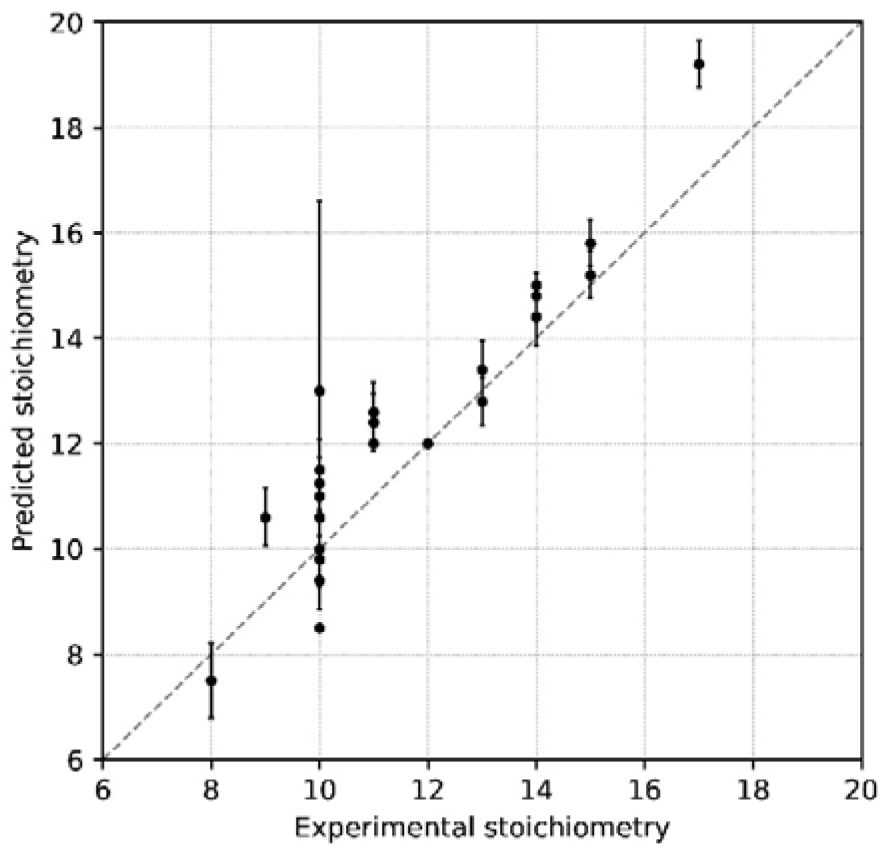
Comparison of predicted stoichiometry values against the experimental data for the set of c-rings with experimentally determined stoichiometries (Supporting Table 1). Means and standard deviations of values obtained from 5 models are shown. Pearson correlation coefficient is equal to 0.94.

To further test the accuracy of the algorithm, we evaluated its predictions for a set of artificially mutated c subunits (Supporting Table 3). While predicted stoichiometry for wild type c-rings did not always match the experimentally observed one, the difference between experimentally obtained stoichiometries of wild type and mutant c-rings led to changes in the stoichiometries predicted by the algorithm in five out of six cases. At the same time, none of the three mutations that preserved stoichiometries of c-rings did not lead to changes in the predicted stoichiometries. This result is even more notable given that evolutionary information used by AlphaFold2 could bias the predicted structures and stoichiometries towards those of natural proteins; predicting effects of mutations using AlphaFold2 is not straightforward (Buel and Walters 2022; McBride et al. 2023; Pak et al. 2023).

### Predicted diversity of natural c-rings

To evaluate the diversity of natural c-rings, we extracted the amino acid sequences from the InterPro database annotated to belong to the family IPR000454 (“ATP synthase, F0 complex, subunit C”; spelling matches that in the database). We filtered out the sequences that met at least one of the following conditions: 1) length is more than 250 amino acids; 2) undefined amino acids present; 3) at least two out of four conserved amino acids (RQPD, RQPE, KQPE or RNPS (Pogoryelov et al. 2012) in the hairpin loop between the c subunit transmembrane helices are not found. Consequently, we obtained a list of 58,732 sequences. Using sequence identity threshold of 60%, the sequences were grouped into 1,571 clusters, and one representative protein from each cluster was modeled and analyzed.

Stoichiometry values predicted for these sequences ranged from 4 to 27, with several cases above 30. Upon closer examination, it turned out that all models with a stoichiometry of less than eight corresponded to unreasonable arrangements, with the possible exception of two sequences, where, out of five models, 1 or 2 corresponded to stoichiometry of 7, and the rest were symmetric dimers (UniProt IDs C3YL44 and A0A221SD96, respectively). Models with stoichiometries above 30 mostly came from unannotated metagenomic data and could not be reasonably verified; raw data is available as Supporting information. At the same time, we obtained a large number of adequate models with the predicted stoichiometry non-negligibly higher than the reported maximum of 17 for F-type ATP synthase (Figure 3A). Most of the obtained models are circular and not helix-like (Figure 1B,C), as reflected in the distribution of the RMSD values for the first and the next-after-the-last protomers for the modeled assemblies (Figure 3B). The examples of constructed c-rings in the whole range of stoichiometries are shown in Figure 3C.

**Figure 3.**
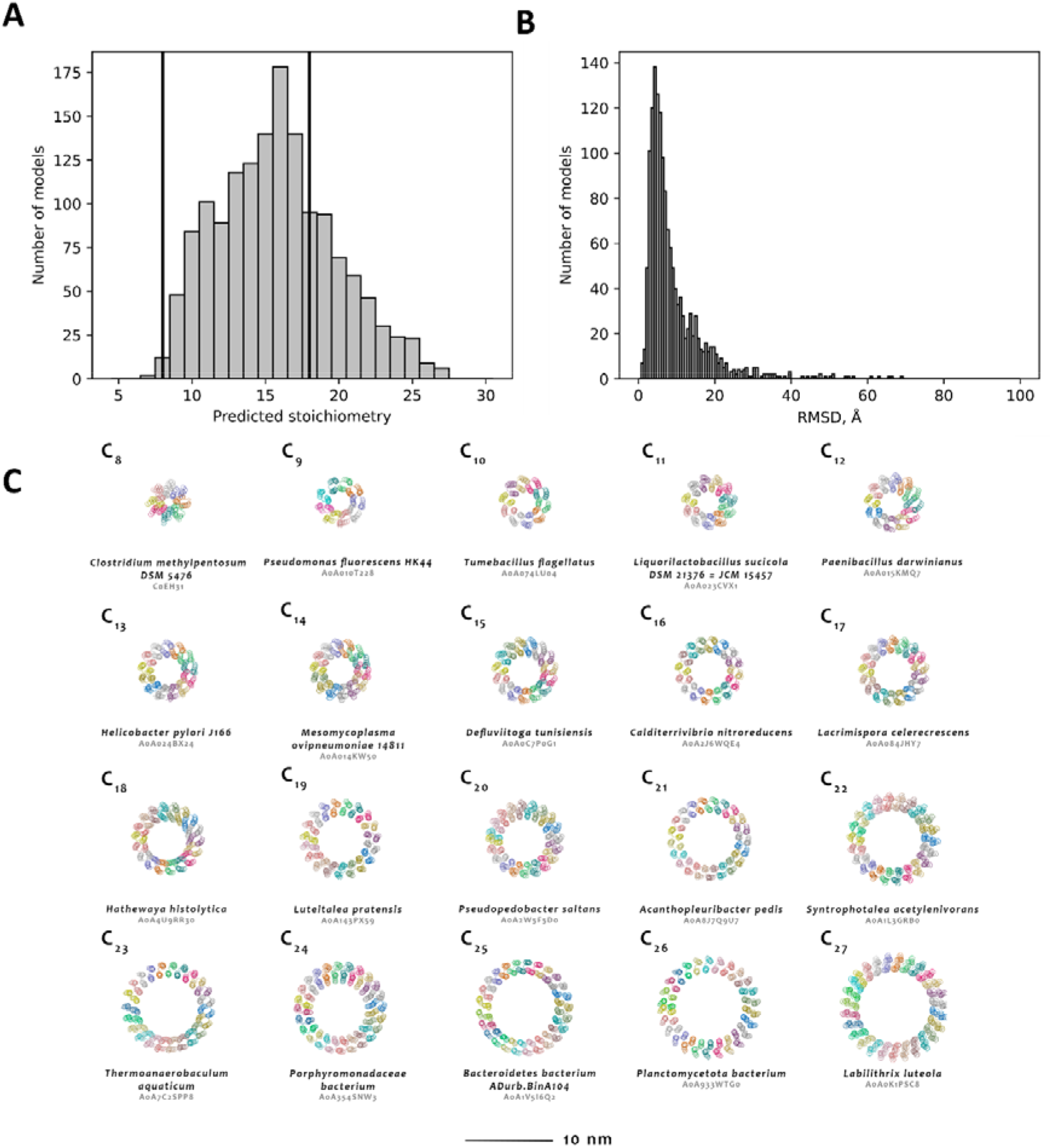
Predicted diversity of natural c-rings. **(A)** Distribution of predicted stoichiometries for representatives of ∼1,500 clusters. Bold vertical lines mark the minimum and maximum stoichiometries that have been experimentally observed. Significant part of the predicted stoichiometries lies outside of this range. In particular, one sequence has the predicted stoichiometry of 7, and 508 sequences of more than 17. **(B)**Distribution of RMSDs of heavy atom positions between the first and the next-after-the-last protomers for the obtained models. In most cases, the ring closes almost perfectly, yet in others there is a notable helical pitch (as shown in Figure 1B,C). **(C)** Examples of predicted c-ring models with stoichiometry ranging from 8 to 27.

As the next step, we built a phylogenetic tree showing predicted stoichiometry for the representative sequences (Figure 4). Strongly related sequences mostly had similar stoichiometries, and multiple occurrences of low and high stoichiometries were observed for different taxonomic groups.

**Figure 4.**
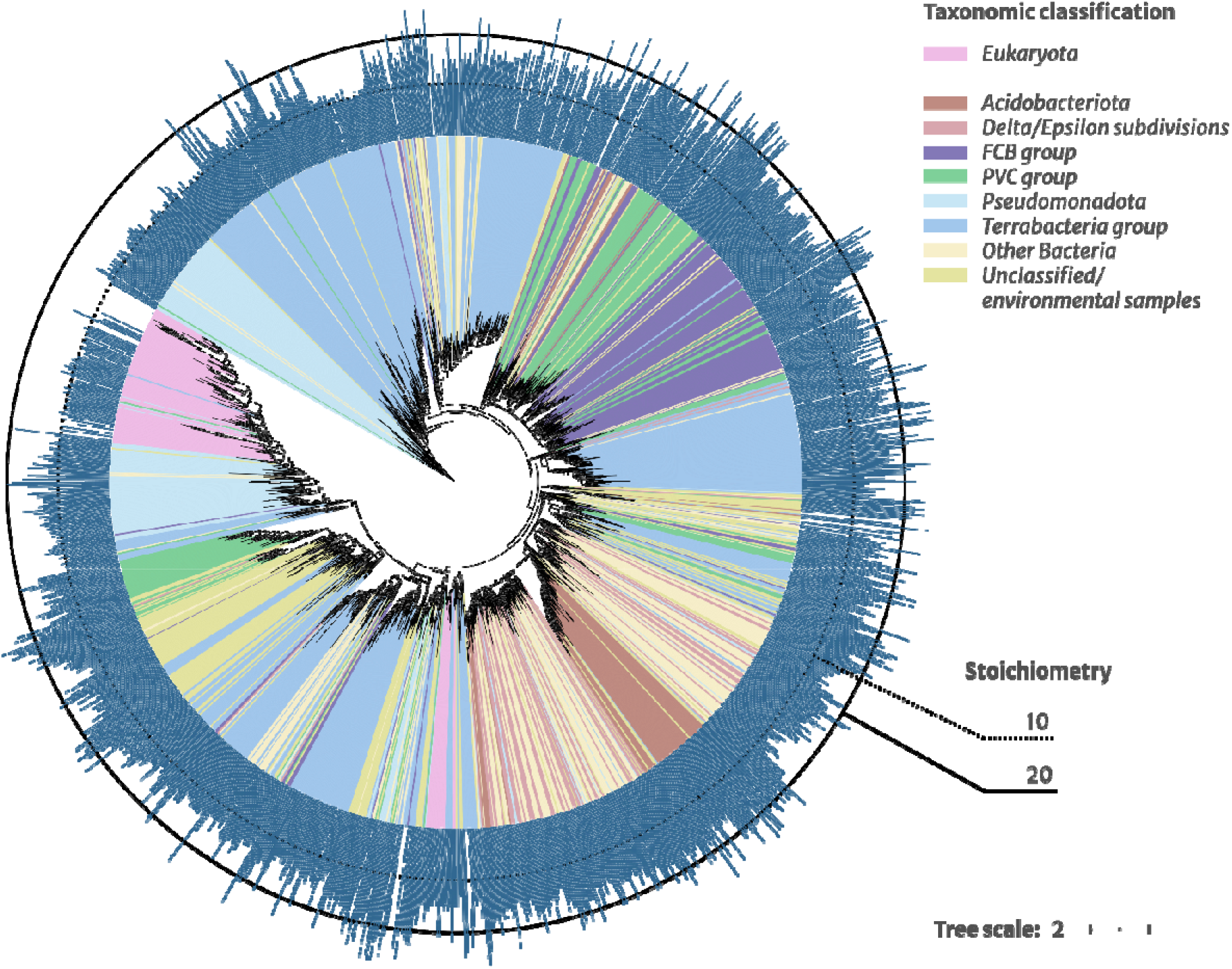
Phylogenetic tree of selected natural c-rings. Sequences are colored by taxonomic classification of the host organisms. Multiple occurrences of low (below 10) and high (above 20) stoichiometries are observed for different taxonomic groups. Blank bars correspond to the cases where stoichiometry was not predicted.

### Molecular dynamics simulations of high-stoichiometry c-rings

Following high-throughput analysis, we wanted to probe whether predicted high stoichiometries are reasonable. We looked for proteins from species with fully sequenced annotated genomes and selected c subunits from *Candidatus Kryptonium thompsoni* (Uniprot ID A0A0P1LIH4, predicted stoichiometry of 23) and *Thalassoglobus polymorphus* (Uniprot ID A0A517QRU4, predicted stoichiometry of 26) for further investigation. We checked and verified that the corresponding operons harbor full sets of ATP synthase subunits (Figure 5) and that the corresponding genomes do not contain other genes encoding c subunits.

**Figure 5.**
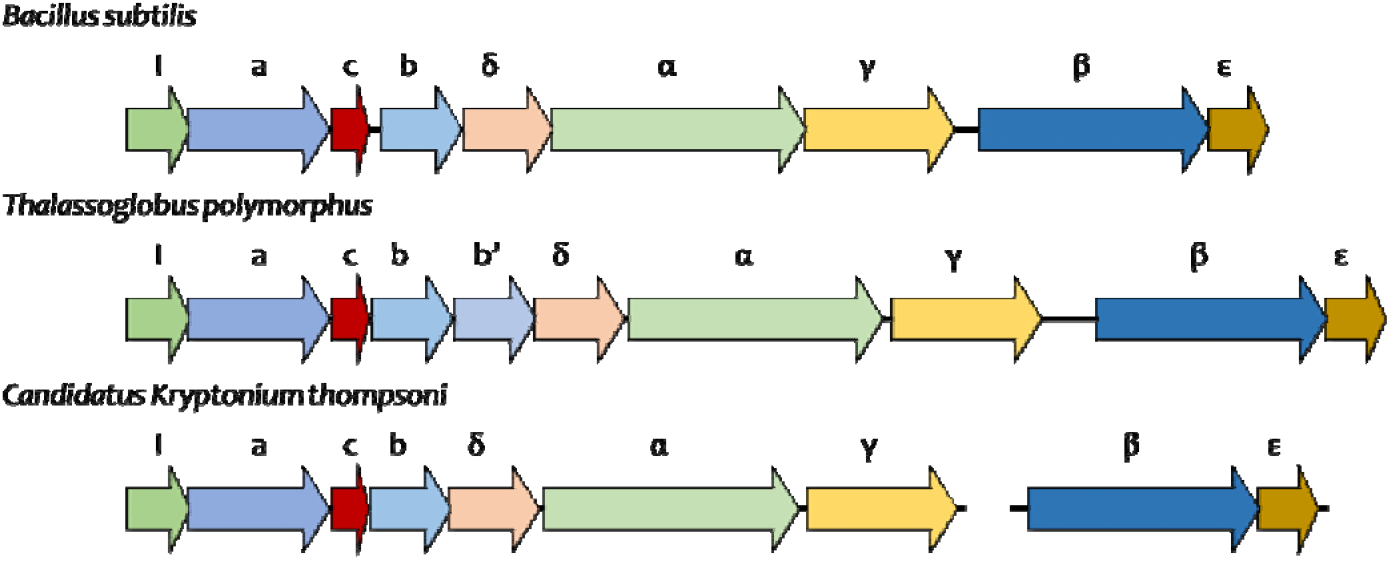
ATP synthase operons from *Bacillus subtilis, Thalassoglobus polymorphus*, and *Candidatus Kryptonium thompsoni*. Operon from *Bacillus subtilis* (strain 168) is used as a reference (Santana *et al*. 1994).

Molecular dynamics (MD) simulations, while requiring accurate execution, currently provide reliable information about behavior of proteins (Hollingsworth and Dror 2018). In comparison with experiments, simulations can be set up relatively fast, without the need for expression and purification of the studied proteins. On the other hand, simulations do not provide a fully definitive answer, because, for example, the effects of the cofactors that are not explicitly included in the simulation will not be taken into account, and in case of c-rings the proteins may be post-translationally modified, or some more-or-less unusual lipids may bind preferentially to the rotor rings. Keeping this and other limitations in mind, we proceeded to model the two selected rotor rings, as well as the spinach chloroplast ATP synthase rotor ring as a control, given that it was well characterized experimentally (Seelert, Dencher and Müller 2003; Balakrishna et al. 2014; Hahn et al. 2018; Vlasov et al. 2019) and studied by us previously using molecular dynamics (Novitskaia, Buslaev and Gushchin 2019).

Because the fully assembled ring may remain stable on the timescale accessible in MD simulations even if the stoichiometry is not correct, we modeled partial assemblies (4, 11, 16 or 18-mers) of subunit c from *Candidatus Kryptonium thompsoni* and *Thalassoglobus polymorphus* in a model bacterial lipid membranes (3:1 POPE:POPG) for 500 ns. We used AlphaFold2 to obtain starting models of the corresponding oligomers in each case, and followed the time evolution of the inter-protomer angles α, with the assumption that the equilibrium value of α will correspond to stoichiometry as follows: N = 360º/α. Irrespective of the inter-protomer angles in the starting models, in each case the angles in each simulation eventually converged to roughly the same values, corresponding to stoichiometries of 21-22 for *Candidatus Kryptonium thompsoni* and 24-25 for *Thalassoglobus polymorphus* (Figure 6 and Supporting Figure 3). Control simulations, where the starting configurations were derived from AlphaFold2 models with deliberately wrong inter-protomer angles, converged to similar values (Supporting Figure 4), including the stoichiometry of around 16 for spinach chloroplast ATP synthase rotor ring (experimentally determined stoichiometry is 14). Thus, we conclude that there is a general correspondence between the AlphaFold2-derived and MD-derived stoichiometries, and presence of unusually high stoichiometries among the natural rotor rings is highly likely.

**Figure 6.**
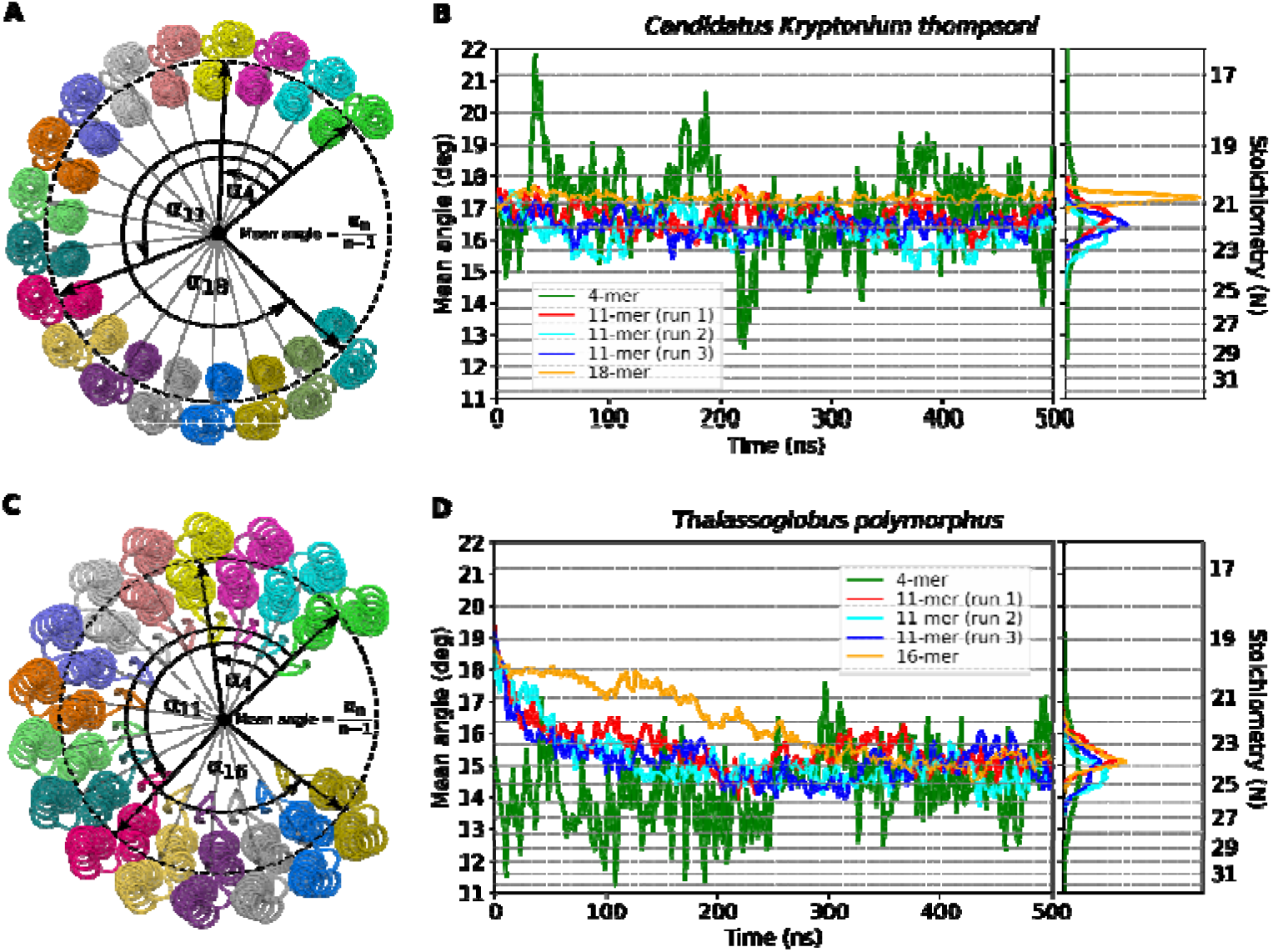
Molecular dynamics simulations of predicted c-rings from *Candidatus Kryptonium thompsoni* and *Thalassoglobus polymorphus*. We modeled incomplete rings to allow for conformational changes. (**A, C**) Employed definitions of mean angles. (**B, D**) Time evolution of mean interprotomer angles for 4, 11, 16 or 18-mers of respective partial rings. Data for each particular angle may be found in Supporting Figure 3. Fluctuations are diminished after 250 ns, presumably due to equilibration of modeled systems; consequently, distributions are calculated for last 250 ns of each simulation. Corresponding stoichiometries are unusually large (clearly above 17) and consistent with AlphaFold2-based predictions. For each simulation, starting oligomer model was independently generated using AlphaFold.

## Discussion

In recent years, machine learning-based approaches revolutionized structural biology providing unexpected advances in structure prediction and protein engineering (Anishchenko et al. 2021; Baek et al. 2021; Jumper et al. 2021; Lin et al. 2023; Watson et al. 2023). AlphaFold2, and built upon it user-friendly and fast implementation ColabFold allowed exploration of the universe of protein structures at a scale that was never available before (Mirdita et al. 2022; Barrio-Hernandez et al. 2023). While originally such approaches were designed to primarily predict structures of isolated single proteins, it was quickly discovered that they also work well for many types of protein complexes (Humphreys et al. 2021; Richard et al. 2021; Akdel et al. 2022; Bryant, Pozzati and Elofsson 2022). More elaborate approaches were later developed to allow for modeling large multi-subunit homo- or heterooligomeric complexes (Bryant et al. 2022; Jeppesen and André 2023; Shor and Schneidman-Duhovny 2024; Schweke et al. 2024).

Here, we present a minimalistic AlphaFold2-based approach for prediction of stoichiometry of homooligomeric complexes, primarily ATP synthase rotor rings, where the model of a dimer is extrapolated to obtain a circular assembly (Figure 1). The approach works well resulting in high correlation between predicted and experimentally determined stoichiometries for test cases (Figure 2), while highlighting that simple sequence-similarity based modeling would not be enough to obtain a good model for most c-rings (Figure 4). The approach tends to slightly overestimate the stoichiometry in some cases, both compared to experimental data (Figure 2) and to results of molecular dynamics simulations (Figure 6); since it is not fully clear how to take this overestimation into account correctly, we did not include any explicit corrections in the approach. Interestingly, using trimers or tetramers as the templates for obtaining the complete assembly does not improve the accuracy of the predictions (Supporting Table 2). We also observe that the algorithm is capable of evaluating the effects of mutations (Supporting Table 3), which is even more remarkable given that no atomistic structures for mutants are available, and consequently the structures were not included in the train set during AlphaFold2 development. In other cases, prediction of the effects of mutations using AlphaFold2 required significant elaboration (Buel and Walters 2022; McBride et al. 2023; Pak et al. 2023). We note that ATPase rotor rings can be considered a special case of coiled coils (Lupas and Bassler 2017; Woolfson 2023), and AlphaFold2 was already used to identify new types of such structures in putative proteins found in sequence databases (Martinez-Goikoetxea and Lupas 2023).

On one hand, our results highlight the advantages of AlphaFold2 and ColabFold. On the other, they also illustrate some of the nuances of using machine learning-based algorithms. Indeed, in a particular example, when prompted, AlphaFold2 generates circular assemblies for a very wide range of stoichiometries (10 to 21) of spinach chloroplast ATP synthase rotor ring, which is well known to be composed of 14 subunits (Supporting Figure 1). Presumably, the algorithm prefers to match all possible interfaces and/or leave no exposed interfaces, generating an incomplete ring only when the stoichiometry is very low (9) or two complete rings when the provided stoichiometry is high enough (22). Also, apparently, AlphaFold2 is able to produce small and large, partial and complete assemblies of transmembrane rotor rings without any explicit knowledge about lipid membranes in general.

Regarding the approach presented in this work, it also inevitably has some limitations. First, the approach relies on obtaining a circular assembly; for most membrane proteins, it is relatively clear that they should fit into the (mostly) planar membrane, yet for soluble proteins some other considerations would be needed to clarify whether the protein forms circular or helical assemblies, and which period and pitch might have helical assemblies (as discussed for example by Boyer et al. 2015). The approach also explicitly relies on the assumption that the assembly is homooligomeric. Second, the structure of the proteins, and in particular, rotor rings (Nesci *et al*. 2016), may be affected by post-translational modifications, which may affect the complex formation but are not considered here. Third, different organisms have different membrane lipid compositions, in general, and in different conditions (Harayama and Riezman 2018); some lipids bind preferentially to rotor rings, and thus might affect the assembly properties. Finally, while using dimers as templates for obtaining complete assemblies is very computationally efficient, slower advanced approaches (Bryant et al. 2022; Schweke et al. 2024; Shor and Schneidman-Duhovny 2024) might have produced even better results.

As for ATP synthases, they are widely studied due to their key role in every living organism. At the same time, being multisubunit transmembrane complexes, they are difficult objects for structural biologists. While tens of thousands of genomes have been sequenced due to advances in genomics, only slightly above 30 rotor rings of F-Type ATP synthases have been structurally characterized (Supporting Table 1). Therefore, we attempted to evaluate the natural diversity of the rotor rings using genomic data and high-throughput stoichiometry prediction. While we didn’t observe robust evidence for rotor rings with stoichiometry below that observed in experiments (8), we found many cases with stoichiometry notably above what was observed (Figures 3, 4). For two selected examples (Figure 5), molecular dynamics simulations (Figure 6) corroborate the AlphaFold2-based predictions. Thus, we conclude that it is highly likely that some natural organisms harbor F-Type ATP synthases with as of yet unexplored stoichiometries. At the same time, given that we used high-throughput approach and didn’t screen manually all of the predicted models, there is a possibility of inadvertent errors in the dataset, and each particular case has to be analyzed carefully before making further assumptions based on it. Eventually, experimental validation is clearly needed to verify these predictions. In any case, the algorithm provides an exciting opportunity of using it to better study the evolution of ATP synthases across the tree of life (Mahendrarajah *et al*. 2023) and to predict the effects of mutations on stoichiometry and engineer ATP synthases tailored to specific applications (Pogoryelov et al. 2012; Preiss et al. 2013; Yamamoto et al. 2023).

High predicted stoichiometries of some rotor rings predicted here raise a number of questions. First, the details of interactions between the soluble and the membrane parts in such ATP synthases are unclear. Namely, the gamma and epsilon subunits interact with the conservative interhelical loops of c-ring subunits. The number of c protomers involved in contacts with epsilon or gamma subunits varies in different organisms. For example, 6 out of 8 c protomers contact the F_1_ part of mitochondrial ATP synthase from *Bos taurus*; 10 out of 14 c protomers contact the F_1_ part of chloroplast ATP synthase from *Spinacia oleracea*; and 5 out of 10 c protomers contact the F_1_ part of bacterial ATP synthase from *Escherichia coli*. If the number of c subunits is above 20, much of the ring will not interact with other subunits assuming standard architecture of the complex.

Another question concerns the interior of rotor rings (discussed in Vlasov *et al*. 2019, Novitskaia, Buslaev and Gushchin 2019). Different possibilities of what constitutes the “plugs” are described or discussed in the literature: lipids (Meier et al. 2001; Novitskaia, Buslaev and Gushchin 2019), protein subunits (Gu *et al*. 2019) or even quinones (Vlasov et al. 2019). In case of c-rings with high stoichiometries, the inner space is large: the inner diameter of the predicted c_27_-ring from *Labilithrix luteola* is about 7 nm (see Figure 3). Such rotor rings may capture many different membrane molecules, and their structural organization and dynamics is unclear.

Finally, one of the most important implications of high c ring stoichiometry is unusual cellular bioenergetics of the respective organism, given that one complete turn of the rotor results in synthesis of 3 ATP molecules. Thus, either such high stoichiometry ATP synthases function at a relatively low electrochemical potential gradient (Preiss *et al*. 2013), or they are comparably inefficient, spending more energy per turn when needed. Previously, high stoichiometry was documented to be useful for alkaliphilic extremophiles (Preiss *et al*. 2013) and photosynthetic organisms (Davis and Kramer 2020; Cheuk and Meier 2021).

## Conclusions

In this work, we presented a simple high-throughput approach for rapid evaluation of stoichiometry of homooligomeric assemblies that allowed us to characterize natural diversity of F-Type ATP synthase rotor rings. We predict that some natural organisms harbor ATP synthases with rotor rings, whose stoichiometry is above what was previously observed in experiments, and which might operate at lower-than-expected transmembrane potentials or higher adverse ion concentration gradients. The approach could likely be used to study other families of proteins that form high order homooligomers with circular symmetry, as well as for engineering protein complexes with altered stoichiometry. Our results further highlight the utility of AlphaFold2, and machine learning-based methods in general, for structural biology applications.

## Materials and Methods

### Preparation of dimer models and prediction of stoichiometry

The models of dimers of c subunits were obtained using ColabFold v1.5.3 (Mirdita *et al*. 2022b). Modeling parameters are presented in Table 1. For each model, we searched a conservative loop in the sequence by aligning the sequence to that from *E. coli* and determined two α-helices connected by the loop. If the model contained any other residues beyond those of *E. coli* sequence, or in non-α-helical conformation, they were removed. The stoichiometry was estimated using the approach described in Figure 1. The script used to obtain stoichiometry based on a model of a dimer is available as a Supporting Information file.

### Analysis and clustering of natural c subunit sequences

Original sequences were downloaded from InterPro on 07.07.2023. Outlier sequences were removed as described in the main text. Remaining 58,732 sequences were clustered using cd-hit v. 4.8.1 (Li and Godzik 2006), with the minimum sequence identity of 60% within each cluster. According to the selected parameters, the sequences were combined into 1,571 clusters, and one sequence was selected from each cluster for further evaluation.

### Phylogenetic tree

Multiple sequence alignment was performed using the MUSCLE algorithm (Edgar 2004). Obtained alignment was trimmed around the edges of *E. coli* ATP synthase subunit c (Genbank GI 56404993). The phylogenetic tree of ATP synthase subunit c sequences was calculated using IQ-TREE v. 1.6.12 (Minh *et al*. 2020). Illustrations of the phylogenetic tree were prepared using iTOL v. 5 (Letunic and Bork 2021). Default sets of parameters were used for all the algorithms. Taxonomy data for each protein were obtained from NCBI Taxonomy browser (https://www.ncbi.nlm.nih.gov/Taxonomy/taxonomyhome.html/index.cgi).

### Molecular Dynamics Simulations

AlphaFold2 4, 11, 16 or 18-mer structures of the subunit c from *Candidatus Kryptonium thompsoni* and *Thalassoglobus polymorphus*, and AlphaFold2 11-mer structure of the subunit c from *Spinacia oleracea* were simulated in a lipid bilayer. Hexagonal unit cells contained native-like bacterial inner membrane lipids POPE/POPG in the proportion 3:1, namely 300 POPE and 100 POPG per monolayer (for *Candidatus Kryptonium thompsoni* and *Thalassoglobus polymorphus*) and 400 POPC lipids per monolayer (for *Spinacia oleracea*). The membranes were solvated with water; 0.15 M NaCl ions were added. All systems were prepared using the Membrane builder tool of CHARMM-GUI web server (Jo *et al*. 2008). Ions were generated using the Distance method of CHARMM-GUI. Protonation states of titratable amino acids were assigned in accordance with pH 7.0 using PROPKA3 (Olsson *et al*. 2011); glutamates and aspartates in the middle of the membrane were protonated.

MD simulations were conducted using GROMACS 2022 (Abraham *et al*. 2015) and the CHARMM36m force field (Best *et al*. 2012) with TIP3P water model (Jorgensen *et al*. 1983). Systems were energy minimized using the steepest descent method, thermalized and equilibrated for 30 ns with restraints on protein backbone atoms, simulated for 300 ns (for *Spinacia oleracea*) and 500 or 600 □ ns (for *Candidatus Kryptonium thompsoni* and *Thalassoglobus polymorphus*) using the leapfrog integrator with a time step of 2 □ fs, at a reference temperature of 303 K and at a reference pressure of 1 □ bar. Temperature was coupled using Nosé-Hoover thermostat(Evans and Holian 1985) with coupling constant of 1 □ ps^−1^. Pressure was coupled with semiisotropic Parrinello-Rahman barostat (Parrinello and Rahman 1981) with relaxation time of 5 □ ps and compressibility of 4.5×10^−5^ □ bar^−1^.

The simulations were performed using periodic boundary conditions. The center of mass of the reference structure was scaled with the scaling matrix of the pressure coupling. The covalent bonds to hydrogens were constrained using the LINCS algorithm (Hess *et al*. 1997). The nonbonded pair list was updated every 20 steps with the cutoff of 1.2 □ nm. Force based switching function with the switching range of 1.0 – 1.2 □ nm and particle mesh Ewald (PME) method (Petersen 1995) with 0.12 □ nm Fourier grid spacing and 1.2□nm cutoff were used for treatment of the Van der Waals and electrostatics interactions.

In-house Python script was used for analysis of the angles between neighboring protomers. The membrane plane was obtained using the Principal Component Analysis for the center of mass of the protomers. Projections of the centers of mass of all protomers onto the membrane plane were fitted with a circle, and each angle was defined as the angle between two vectors connecting the center of the circle with the projections of the centers of mass of the neighboring protomers onto the membrane plane.

## Supporting information

Archive including all Supporting Information data

## Acknowledgments

Literature analysis and evaluation of predicted stoichiometries was supported by the Russian Science Foundation project 22-74-00044. Structure prediction and modeling was supported by the Ministry of Science and Higher Education of the Russian Federation (agreement 075-03-2023-106, project FSMG-2021-0002).

## Supporting information

The following supporting information is available:

File with Supporting Tables 1-3 and Supporting Figures 1-4.

Files with predicted stoichiometry data for experimentally characterized rotor rings, selected sequences, high stoichiometry proteins and all analyzed sequences from InterPro.

Python script used to estimate stoichiometry using a dimer model.

## Data availability

Molecular dynamics trajectories have been deposited to Zenodo and are available using the following link: doi.org/10.5281/zenodo.10670316. All other data is available as Supporting Information.

## Code availability

Python script used to estimate stoichiometry using a dimer model is available as a Supporting Information file.

