## Supplementary material for "High-throughput algorithm predicts F-Type ATP synthase rotor ring stoichiometries of 8 to 27 protomers": Archive including all Supporting Information data: 240227 c-ring stoichiometry SI SO IG.pdf

**Supporting Tables 1-3**

**Supporting Figures 1-4**

**Supporting Table 1.** Predicted stoichiometry data for natural c-rings.

| Species name | Experimental stoichiometry | PDB ID (if available) | Predicted stoichiometry |
|---|---|---|---|
| *Acetobacterium woodii* | $9_F/1_V$ | 4BEM | 12.4 |
| *Arthrospira platensis* | 15 | 2XQS, 2XQT, 2XQU, 2WIE | 15.2 |
| *Bacillus pseudofirmus* of4 | 13 | 4CBJ, 4CBK, 2X2V | 13.4 |
| *Bacillus* sp ps3 | 10 | 6N30; 6N2Z; 6N2Y | 11.5 |
| *Bacillus* sp ta2 a1 | 13 | N/D | 12.8 |
| *Bos taurus* | 8 | 5FIL; 5ARI; 5ARH; 5FIK; 5FIJ; 5ARA; 5ARE | 7.5 |
| *Burkholderia pseudomallei* | 17 | N/D | 19.2 |
| *Clostridium paradoxum* | 11 | N/D | 12 |
| *Escherichia coli* | 10 | 5T4P; 5T4O; 5T4Q | 10.6 |
| *Euglena gracilis* | 10 | 6TDU; 6TE0; 6TDY; 6TDZ | 8.5 |
| *Fusobacterium nucleatum* | 11 | 3ZK1; 3ZK2 | 12.4 |
| *Gloeobacter violaceus* pcc 7421 | 15 | N/D | 15.8 |
| *Ilyobacter tartaricus* | 11 | 2WGM, 1YCE | 12.6 |
| *Mycobacterium smegmatis* | 9 | 7NJK, 7NJL, 7NJM, 7NJN, 7NJO, 7NJP, 7NJQ, 7NJR, 7NJS, 7NJT, 7NJU, 7NJV, 7NJW, 7NJX, 7NJY, 7NK7, 7NK9, 7NKB, 7NKD, 7NKH, 7NKJ, 7NKK, 7NKL, 7NKN, 7NKP, 7NKQ, 7NL9 | 10.6 |
| *Mycolicibacterium phlei* | 9 | 4V1F, 4V1G, 4V1H | 10.6 |
| *Nicotiana tobacum* | 14 | N/D | 14.8 |
| *Ogataea angusta* | 10 | N/D | 10.6 |
| *Paracoccus denitrificans* | 12 | N/D | 12 |
| *Paramecium tetraurelia* | ~10 | N/D | 10 |

| | | | |
|---|---|---|---|
| *Pisum sativum* | 14 | 3V3C | 15 |
| *Polytomella* sp. | 10 | 6RDV; 6RDE; 6RE4; 6RE1; 6RE7; 6REU; 6RDP; 6REA; 6RDS; 6RDY; 6RED; 6RDJ; 6RDB; 6RDG; 6RDM; 6RER; 6RD9; 6RE2; 6RE3; 6RE5; 6RE6; 6RE0; 6RDX; 6RDW; 6RET; 6RES; 6REB; 6REC; 6RDC; 6RDI; 6RDH; 6REP; 6RDK; 6REF; 6REE; 6RDZ; 6RDR; 6RDL; 6RDU; 6RDT; 6RE9; 6RE8; 6RDO | 13 |
| *Propionigenium modestum* | 11 | N/D | 12.6 |
| *Saccharomyces cerevisae* | 10 | 5BPS, 5BQ6, 5BQA, 5BQJ, 4F4S, 3UD0, 3U2F, 3U2Y, 3U32 | 11 |
| *Spinacia oleracea* | 14 | 6FKI; 6FKF; 6FKH | 15 |
| *Sus scrofa* | 8 | 6J5K; 6J5J; 6J5I | 7.5 |
| *Synechococcus elongatus* pcc 6301 | 14 | N/D | 15 |
| *Synechococcus elongatus* pcc 7942 | 14 | N/D | 15 |
| *Synechocystis* sp pcc 6803 | 14 | N/D | 14.4 |
| *Tetrahymena thermofilia* | 10 | 6YNW | 9.4 |
| *Toxoplasma gondii* | 10 | 6TMG, 6TMH, 6TMI, 6TMJ, 6TMK, 6TML | 11.25 |
| *Yarrowia lipolytica* | 10 | 5FL7 | 9.8 |

Stoichiometry values are obtained from Flygaard et al. 2020; Montgomery et al. 2021; Mühleip et al. 2021; Vlasov et al. 2022; Yamamoto et al. 2023.

**Supporting Table 2.** Pearson correlation coefficients between the experimental and predicted stoichiometries of natural c-rings. AlphaFold2 can generate and rank several models; correlation values calculated for stoichiometry obtained from 1 best model and for average stoichiometry obtained from all 5 models are presented.

| **Modeled oligomer** | **Interface** | **Using 1 best model** | **Using 5 models** |
|---|---|---|---|
| Dimer | #1-#2 | 0.86 | 0.94 |
| Trimer | #1-#2 | 0.65 | 0.90 |
| Trimer | #2-#3 | 0.89 | 0.92 |
| Tetramer | #1-#2 | 0.96 | 0.95 |
| Tetramer | #2-#3 | 0.96 | 0.92 |
| Tetramer | #3-#4 | 0.95 | 0.95 |

**Supporting Table 3.** Predicted stoichiometry data for mutant c-rings.

| Source | Mutation | Experimental stoichiometry | Predicted stoichiometry | Reference |
|---|---|---|---|---|
| *I. tartaricus* | Wild Type | 11 | 12 | Pogoryelov et al. 2012 |
| | G25A | 12 | 12 | |
| | G27A | 12 | 13 | |
| | P28A | 11 | 12 | |
| | G29A | 11-12 | 13 | |
| | Q32A | 11 | 12 | |
| *N. tabacum* | Wild Type | 14 | 15 | Yamamoto et al. 2023 |
| | tobacco-*platensis* (mutant) | 15 | 16 | |
| *B. pseudofirmus OF4* | Wild Type | 13 | 14 | Preiss et al. 2013 |
| | extWT | 13 | 14 | |
| | extA16G | 12-13 | 12 | |
| | extA16/20G | 12 | 13 | |

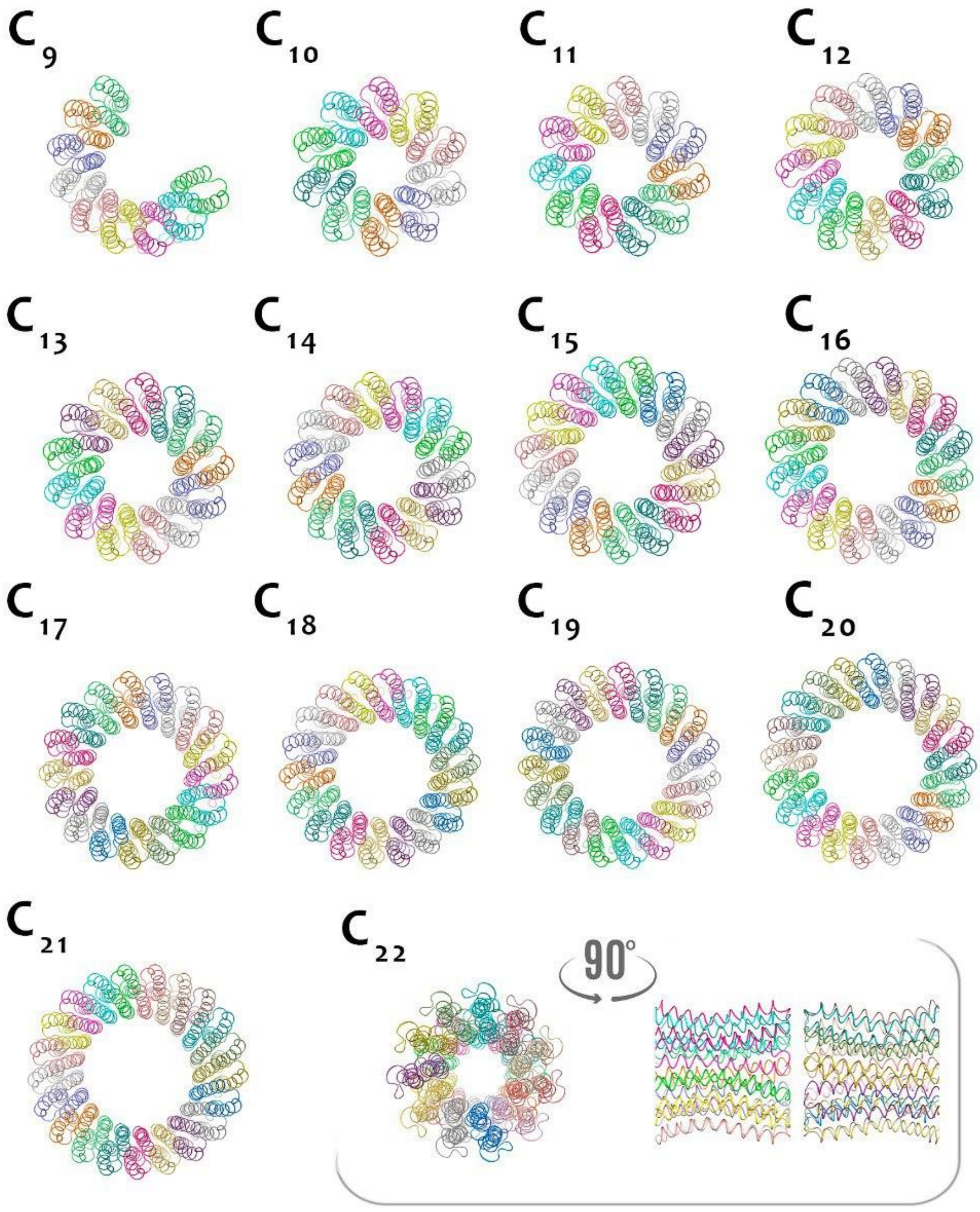


**Supporting Figure 1.** AlphaFold2 models of different oligomers of c subunit of chloroplast ATP synthase from *Spinacia oleracea* with experimentally determined stoichiometry of 14. When predicted using AlphaFold2, 9-mers and below are modeled as incomplete rings (intersubunit arrangement corresponds to 13-mer); 10-mers to 21-mers are modeled as complete rings; and 22-mers are modeled as two stacked 11-mers.

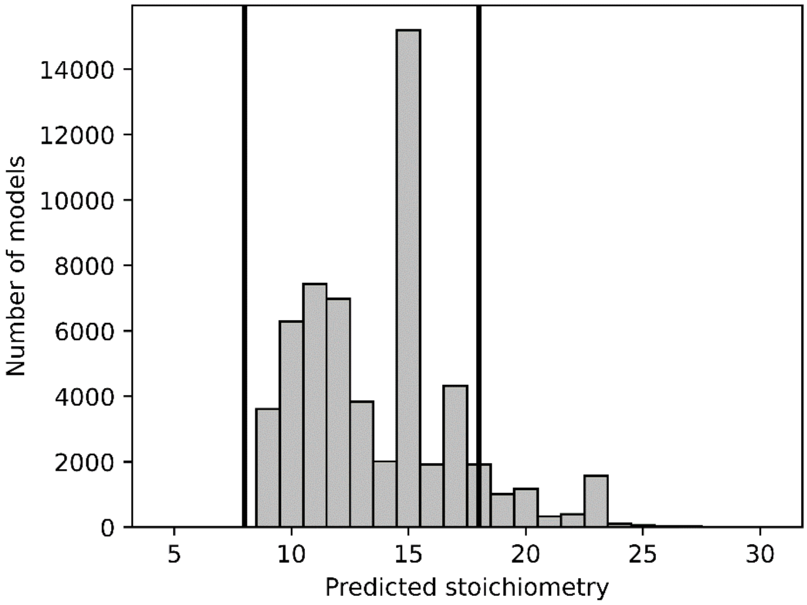


**Supporting Figure 2.** Distribution of predicted stoichiometries corrected for cluster sizes. Stoichiometries were obtained for representatives from ~1,500 clusters of sequences. Bold vertical lines indicate the minimum and maximum stoichiometries that have been experimentally observed (from 8 to 17).

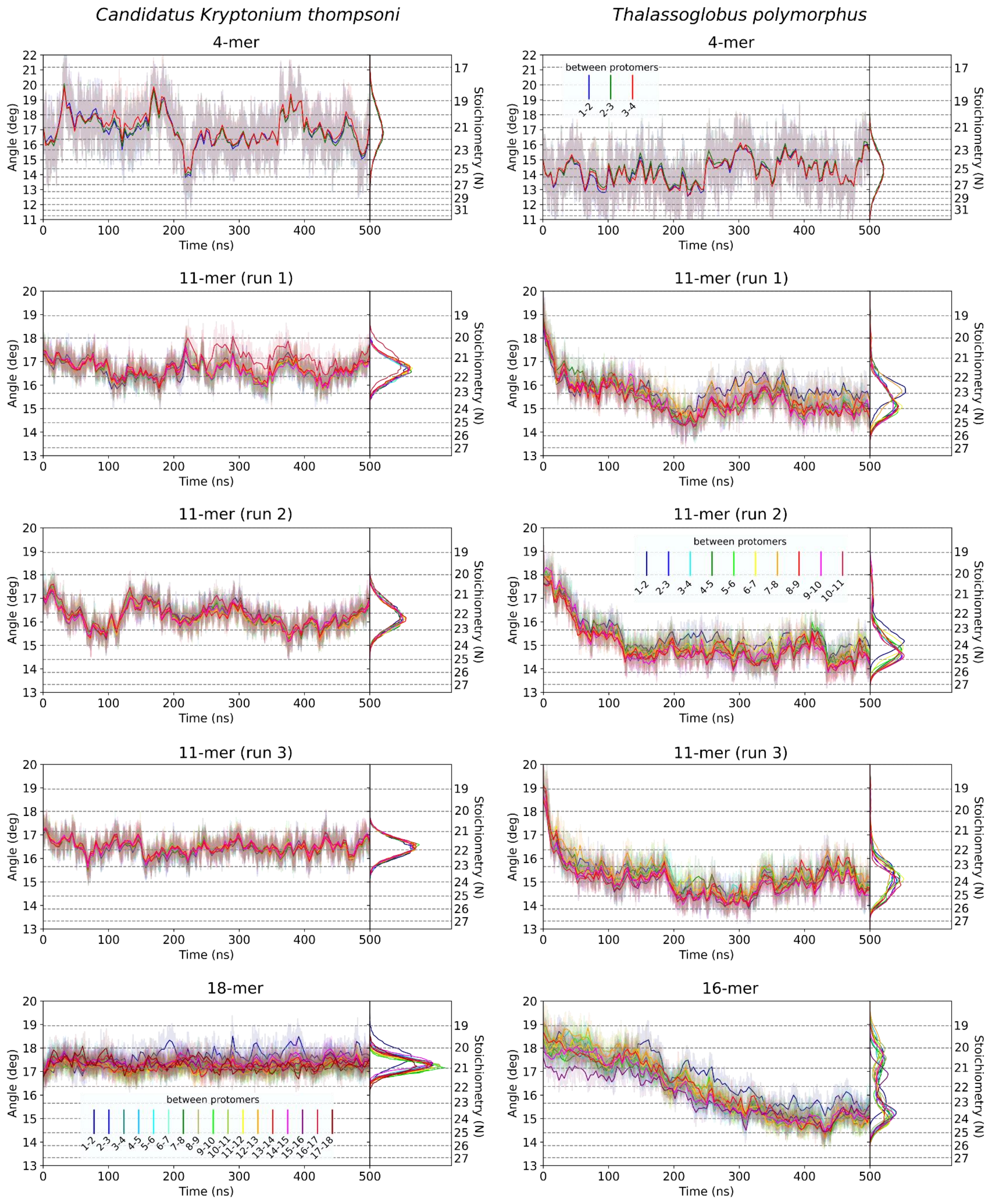


**Supporting Figure 3.** Time evolution and distributions of the angles between neighboring protomers of the AlphaFold2 4, 11, 16 or 18-mers of the subunit c (partial rings) from *Candidatus Kryptonium thompsoni* and *Thalassoglobus polymorphus* in molecular dynamics simulations. The stoichiometry corresponding to these angles is unusually large (above 17) and consistent with AlphaFold2-based predictions.

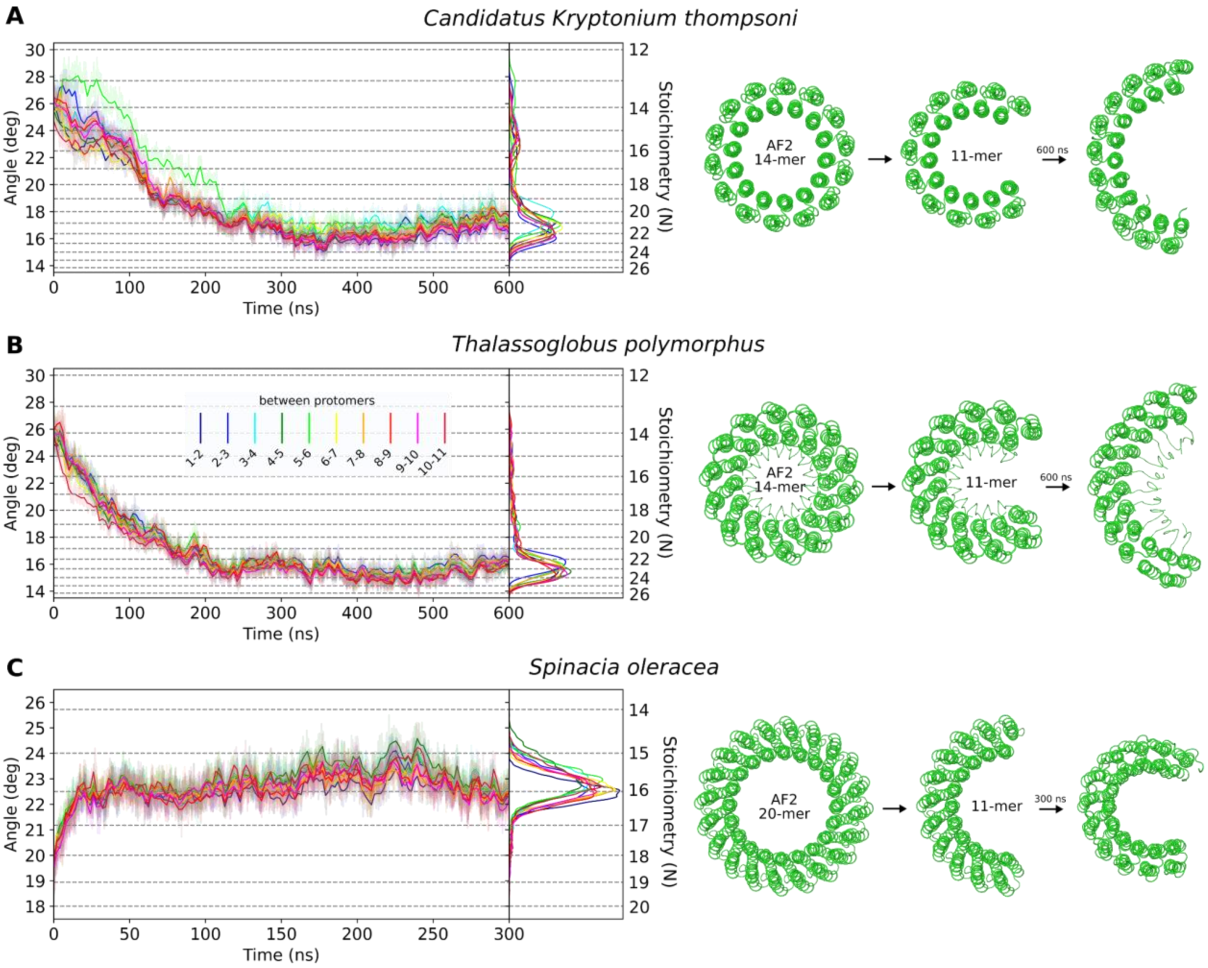


**Supporting Figure 4.** Results of molecular dynamics simulations of c subunit 11-mers using parts of 14- and 20-mers prepared using AlphaFold2 as starting models. In accordance with expectations, time evolution and distributions of the angles between neighboring protomers (left), and final structures from simulations (right) converge towards expected stoichiometries either by extension (*Candidatus Kryptonium thompsoni*, **A**, and *Thalassoglobus polymorphus*, **B**) or by curling (*Spinacia oleracea*, **C**, experimental stoichiometry is 14).
