## Supplementary material for "High-throughput algorithm predicts F-Type ATP synthase rotor ring stoichiometries of 8 to 27 protomers": Archive including all Supporting Information data: Supporting_data_script_stoichiometry_from_dimer.py

```
#python 3.8.8
#pymol 2.4.1
#mdtraj 1.9.6
#matplotlib 3.3.4

#26.02.2024 by Stepan Osipov
#High-throughput algorithm predicts F-Type ATP synthase rotor ring stoichiometry of 8 to 27 protomers
#deposited to biorxiv

from pymol import cmd
import mdtraj as md
import math
import matplotlib.pyplot as plt

def rmsd(chain_1, chain_2, structure):
    N = len(chain_1)
    summ = 0
    for i in range(0, N):
        summ += pow(structure.xyz[0][chain_1[i]][0] - structure.xyz[0][chain_2[i]][0], 2)
        summ += pow(structure.xyz[0][chain_1[i]][1] - structure.xyz[0][chain_2[i]][1], 2)
        summ += pow(structure.xyz[0][chain_1[i]][2] - structure.xyz[0][chain_2[i]][2], 2)
    return math.sqrt(summ/N)*10.0

def rmsd_list_from_dimer(filepath, filename, list_len, temp_file='temporary_file.pdb'):
    cmd.reinitialize()
    cmd.load(filepath + '/' + filename)
    for i in range(0, list_len):
        cmd.copy('c' + str(i), filename[:-4])
    
    for i in range(0, list_len - 1):
        cmd.align ('/c' + str(i + 1) + '//A', '/c' + str(i) + '//B')
    for i in range(0, list_len):
        cmd.delete('/c' + str(i + 1) + '//A')

    cmd.select('all')
    cmd.save(temp_file)
    
    struct = md.load(temp_file)
    chain_list = list()
    for i in range(0, 2*list_len):
        chain = [atom.index for atom in struct.topology.atoms if (atom.residue.chain.index == i)]
        chain_list.append(chain)
    rmsd_list = list()
    for i in range(0, list_len):
        rmsd_list.append(rmsd(chain_list[0], chain_list[2*i], struct))
    return rmsd_list   

def min_rmsd(rmsd_list):
    for i in range(2, len(rmsd_list) - 1):
        if rmsd_list[i] < rmsd_list[i - 1] and rmsd_list[i] < rmsd_list[i + 1]:
            return i

def predict_stoichiometry(filepath, filename):
    return min_rmsd(rmsd_list_from_dimer(filepath, filename, 35))

def draw_rmsd_curve(filepath, filename):
    plt.plot(rmsd_list_from_dimer(filepath, filename, 35))
```
